# Machine and deep learning single-cell segmentation and quantification of multi-dimensional tissue images

**DOI:** 10.1101/790162

**Authors:** Eliot T McKinley, Joseph T Roland, Jeffrey L Franklin, Mary Catherine Macedonia, Paige N Vega, Susie Shin, Robert J Coffey, Ken S Lau

## Abstract

Increasingly, highly multiplexed *in situ* tissue imaging methods are used to profile protein expression at the single-cell level. However, a critical limitation is a lack of robust cell segmentation tools applicable for sections of tissues with a complex architecture and multiple cell types. Using human colorectal adenomas, we present a pipeline for cell segmentation and quantification that utilizes machine learning-based pixel classification to define cellular compartments, a novel method for extending incomplete cell membranes, quantification of antibody staining, and a deep learning-based cell shape descriptor. We envision that this method can be broadly applied to different imaging platforms and tissue types.

## Introduction

Recently, single-cell analytical methods have emerged that are capable of interrogating cellular heterogeneity. These methods allow multiple markers to be measured over a large number of cells and can be achieved by disaggregation of tissue or *in situ* analysis. In the first approach, tissues are separated into single cells that can then be subjected to high throughput single-cell RNA-seq (Tang et al. 2009), multi-channel flow cytometry (Bradford et al. 2004), and mass cytometry (Bendall et al. 2011). However, a major limitation of this approach is that the spatial distribution of cells is lost. *In situ* methods such as isotope-based imaging (Giesen et al. 2014; Angelo et al. 2014) or the various multiplexed fluorescence imaging methods (Gerdes et al. 2013; Lin et al. 2018; Goltsev et al. 2018) retain spatial information but require cell identification after data collection. This, in turn, depends on robust methods to reliably identify cell objects. Cell segmentation can be achieved using a variety of methods *in vitro* (Yin et al. 2010; Zimmer et al. 2002; Al-Kofahi et al. 2018). However, directly applying methods developed *in vitro* to a tissue context is often unreliable. Irregular shapes, high density, membrane polarity, and uneven membrane marker coverage make cell segmentation in tissue particularly challenging. Many tissue segmentation applications identify cell nuclei and subsequent dilation (Schmitt and Hasse 2009) or use Voronoi tessellation or watershed to define a cell (Lin et al. 2018). These methods can work well for segmentation and quantification of immune cells with sparse cytoplasm. However, this type of nuclear dilation does not perform well in other tissue contexts composed largely of polarized epithelial cells often times with elongated shapes. Multiple membrane markers have been used to better define cell borders for seed-based watershed segmentation (McKinley et al. 2017) as well as other methods (Schüffler et al. 2015; Baggett et al. 2005; Santamaria-Pang et al. 2015); however, defining cell membranes, a critical step in overall cell segmentation remains a significant problem.

Here, we demonstrate cell segmentation from multiplexed immunofluorescence-derived images (Gerdes et al. 2013) of colorectal adenomas (Shrubsole et al. 2008). Initial pixel classification by random forest-based machine learning provides an input to a novel method to identify and separate cells with internal membranes. Following cell identification, each maker is then quantified by image intensity over the entire cell as well as in the nucleus, membrane, and cytoplasm. Finally, cell shapes are characterized using a autoencoder neural network.

## Results

Our segmentation pipeline utilizes a combination of Matlab for image processing and quantification and Ilastik (Sommer et al. 2011) for pixel-classification using machine learning. Using Matlab provides flexibility to tailor the segmentation to a particular tissue of interest, while Ilastik provides machine-learning tools for images using a user interface.

A flow chart and visual overview of the pipeline are found in Figure 1.

**Figure 1.**
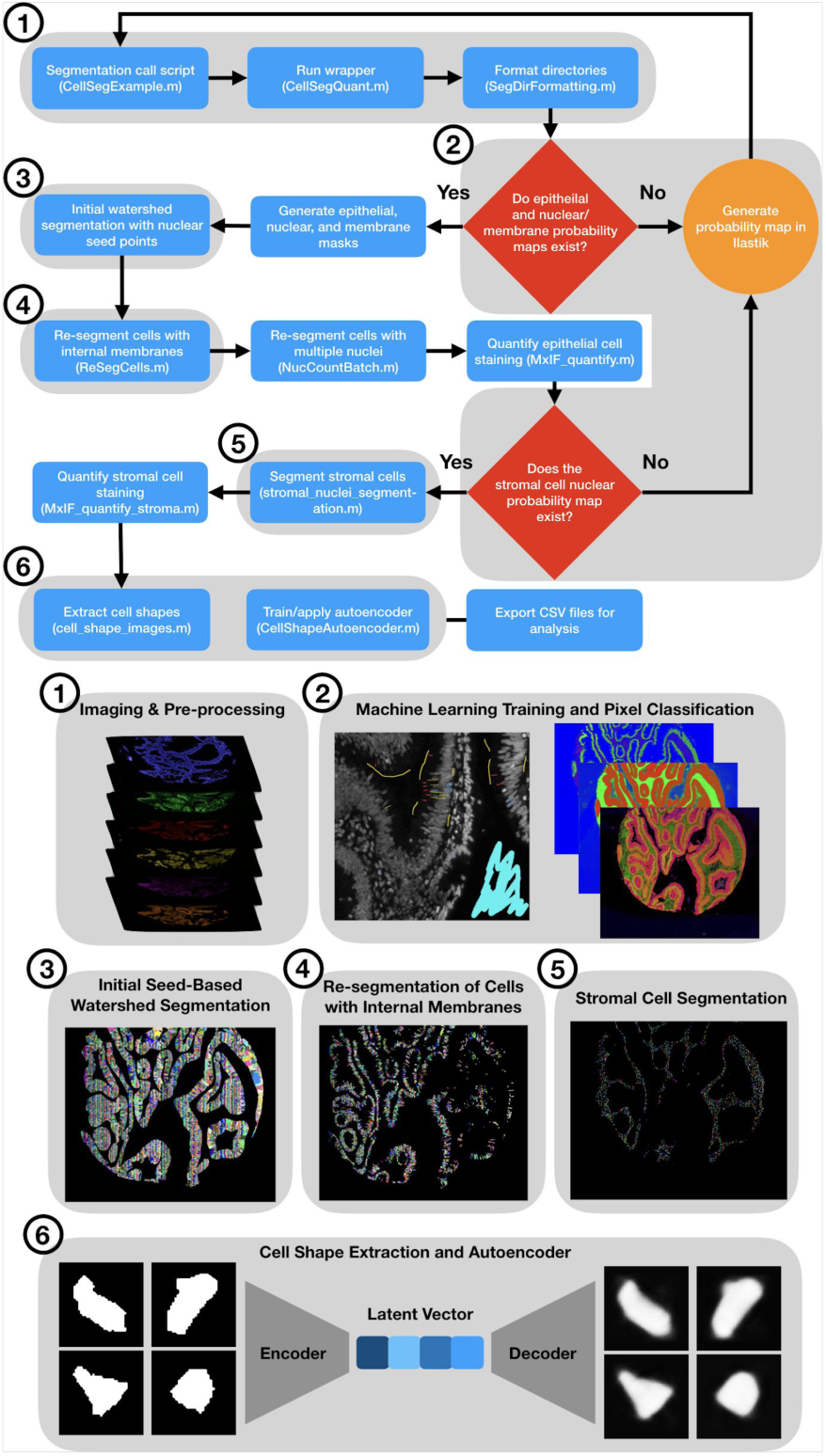
Schematic overview of segmentation pipeline. Flowchart demonstrates the segmentation pipeline steps with corresponding Matlab scripts annotated. Key steps (numbered) are also shown pictorially.

The input to the pipeline can be any type of multiplexed imaging data (e.g. immunofluescence, imaging mass cytometry). Upon initializing the pipeline, various directories are created to store images and tables. Tiff stacks, containing a single marker in each channel, are created in Matlab for processing in Ilastik. The user manually annotates epithelium, stroma, and glass on the tiff stack in Ilastik to generate probability maps for each class. Similarly, the user defines nucleus, cell membrane, cytoplasm, and glass using a separate training session. Optionally, the user can also define epithelial and stromal nuclei if stromal segmentation and quantification are desired. Once the random forest pixel classification steps are completed in Ilastik, the Matlab script is re-initialized.

Masks for the epithelial regions, nuclei, and cell membranes are generated from the probability maps. Subsequently, a watershed is performed on the cell membrane mask using cell nuclei as imposed minimum points to obtain an initial cell segmentation. Due to membrane gaps, this can often result in under-segmented cells, i.e., multiple cells classified as one object. A re-segmentation algorithm is then applied to cells that contain greater than 10% internal membranes by area. This algorithm finds endpoints of internal membranes and extends them until intersecting with either the cell border or another extension from a separate internal membrane. Similarly, in order to maintain intestinal goblet cells, cells with high levels of Muc2 expression are also re-segmented if Muc2 is present in the marker set. Finally, image intensities in the whole cell and subcellular compartments (nucleus, cell membrane, cytoplasm) are quantified.

Stromal cell segmentation is more straightforward due to lower cell density compared to epithelial cells, generally smaller size, and lack of polarity. Stromal cell nuclei are defined by a machine-learning classification and cell segmentation is accomplished by dilation of a set number of pixels from the nucleus. Image intensities are then quantified over the entire cell.

In each case, final cell segmentation maps are produced, with each cell object having a unique pixel value as an identifier. Additionally, sub-cellular localization images are generated to more easily assess the segmentation results. Finally, an autoencoder neural network is trained on binary images of cell shapes derived from single-cell segmentation in the epithelium. Each cell is associated with the encoded latent vector for each cell for downstream analysis. The results of cellular quantification and shape analysis are written as comma separated value (CSV) files. Each row represents a single cell with a distinct identification number and analytes are represented in each column. The CSV files can be used for further analysis using any platform (e.g., R, Python).

Example of raw images, probability masks, and final segmentation are shown in Figure 2. All staining rounds, including DAPI (Fig. 2A) and NAKATPase (Fig. 2B) were used to derive the epithelial (Fig. 2C) and membrane/nucleus (Fig. 2D) probability maps. A final cell segmentation mask is shown in Fig. 2E and sub-cellular segmentation (Fig. 2F) shows the spatial extent of each cell border (white), nucleus (blue), cell membrane (red), and cytoplasm (green). Voronoi segmentation (red) from each nucleus (blue) shows agreement to our segmentation algorithm (green) in areas of high nuclear density (Supplemental Fig. 1). However, when no nucleus is present, Voronoi segmentation, which does not take into account cell membranes (white), fails whereas our algorithm is able to define cells in these regions.

**Figure 2.**
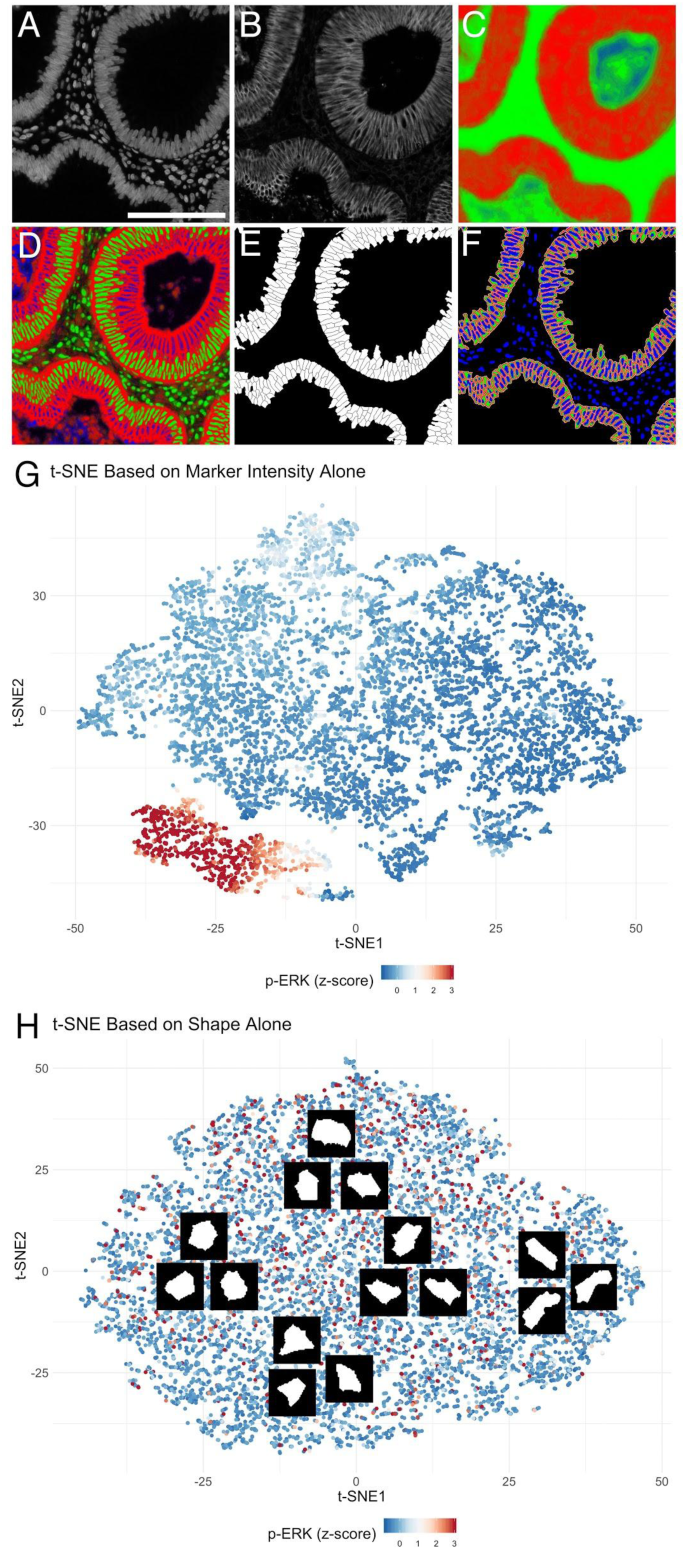
Segmentation pipeline results. An example of selected steps in the segmentation pipeline are shown (scale bar: 100 μm. (A) DAPI and (B) autofluorescence removed NaKATPase images were part of the 20 dimensional image stack used for machine learning. Probability maps for (C) epithelium (red) and stroma (green), (D) cellular membrane (red), nucleus (green), and cytoplasm (blue), as well as (E) final cell segmentation and (F) subcellular segmentation are shown. (G) Results of t-SNE using only marker intensity show distinct populations, such as those marked by p-ERK. However, (H) t-SNE using only the cell shape latent vectors derived from the autoencoder can not segregate cells into populations. Cells of qualitatively similar shape cluster together as shown in the subsetted cell segmentation images.

The CSV files generated from the segmentation pipeline can be analyzed using any number of algorithms, for example, to define populations (Maaten and Hinton 2008) or interrogate cell state transition trajectories (Herring et al. 2018). Figure 2G shows the results of t-stochastic neighbor embedding (t-SNE) using only the intensities for each marker in the data set. Cells with similar expression patterns group together in two-dimensional data space derived from the 18 dimensional data (Supplemental Fig. 2). For example, cells with p-ERK staining intensity form a discrete cluster that not surprisingly overlaps with p-EGFR, PCNA, and Ki67, all of which are features of proliferative cells. When only the encoded latent vector for cell shape is used as an input to t-SNE analysis (Fig. 2H), cells with similar geometries are grouped close to each other.

However, no discernable populations are isolated from the t-SNE analysis, nor are any patterns in marker expression (Supplemental Fig. 3). While cells of different shapes (e.g., columnar, circular, triangular) are able to be identified, and nearest neighbors confirm the shape groupings (Supplemental Fig. 4) cell shape does not appear to correspond to marker signal intensity, at least in this setting.

## Discussion

Cell segmentation represents a difficult task in tissue sections and robust and accurate methods are needed to leverage the rapidly expanding number of highly multiplexed imaging platforms. Furthermore, the diversity of cellular architecture necessitates flexibility in the choice of markers and analytical methods. Our segmentation pipeline allows for this flexibility, and is, in principle, agnostic to the platform used for data collection.

Multiple membrane marker staining intensities have been used to better define the entirety of the cell membrane to aid cell delineation. However, these additive methods suffer from the requirement that only a positive signal from a marker is used to define the cell membrane. A machine-learning pixel classification strategy, like we employ using Ilastik, uses the totality of imaging data to define cell edges not only by intensity but also by edges and image texture. The machine-learning method is better able to capture the entire cell membrane, as well as other cell features such as nuclei and cytoplasm, and is applicable to any tissue type. Similar advantages are obtained using machine-learning pixel classification to define epithelial and stromal masks. This method also requires well-annotated training sets and is more time consuming than additive intensity methods. However, typically the most time-consuming step of multiplexed imaging methods is downstream analysis rather than image collection, processing, or segmentation, so the additional time spent for machine-learning training and application is relatively minor.

Despite improved membrane delineation from machine learning, gaps in the “learned membranes” remain. This is a particular problem in thin sections where a cellular cross section may include any part of the cell. Oftentimes, the cross section may exclude the nucleus, which can result in under-segmentation, or a combined cell with no nucleus to one in which a nucleus is detected. To ameliorate this issue, we developed an algorithm that detects objects in the initial watershed segmentation that have internal membranes. For those cells, the algorithm computes the angle of membrane segments within the cell and extends them until another membrane is encountered. This procedure improves segmentation quality in cases where no nucleus is detected in a neighboring cell.

Additionally, we introduce a deep learning model for classifying cell shape. We chose an autoencoder as other methods that use defined shape descriptors (Möller et al. 2017), such as circularity and concavity may not capture the relevant characteristics that a neural network can. In principle, cell shape should provide added information about cell characteristics, e.g., crypt base columnar cells have an elongated wedge shape and intestinal goblet cells have their eponymous shape. However, when taken in an unknown cross section, this is not always the case. As our results in human colonic adenomas show, 2-dimensional cell shape may not correlate with cell type or protein expression as might be expected if the full 3-dimensional cell morphology was characterized. While this is the case in our system, cell shape may be more informative in other tissue contexts or when limited to already defined cell types.

Together, these improvements to cell segmentation and quantification present a step forward for single-cell analysis of highly multiplexed imaging data. Importantly, these methods are tissue-type and acquisition method independent and are highly modifiable. We envision this pipeline will be broadly applicable to the growing multiplexed imaging community.

## Acknowledgments

Special thanks to Benoit Pimpaud for help developing the shape characterization algorithms. Research reported in this publication was supported by the National Cancer Institute (NIH) under awards R35CA197570, P50CA236733, and U2CCA233291 to RJC, R01DK103831 to KSL. PNV was supported by a training grant funded by NIH under award T32HD007502. The content is solely the responsibility of the authors and does not necessarily represent the official views of the NIH.

## Supplemental Methods

### Human Subjects

The Tennessee Colon Polyp Study (TCPS) (Shrubsole et al. 2008) was approved by the Vanderbilt University Medical Center (VUMC) and Veterans Affairs Tennessee Valley Health System (VA) institutional review boards and the VA Research and Development Committee. All participants provided written informed consent.

### Tissue Processing, Antibody Staining, and Tissue Imaging

A de-identified colonic adenoma tissue microarray (TMA) derived from the TCPS patient cohort was obtained. The TMA was sectioned (5 µm) prior to deparaffinization, rehydrations and antigen retrieval using pH 6.0 citrate buffer (DAKO) at 105°C for 20 minutes followed by 10 minutes at room temperature. The slide was incubated in 3% H2O2 for 10 minutes to reduce endogenous background signal and subsequently blocked in 3% BSA/10% donkey serum in PBS for 30 minutes. Sequential antibody staining and dye inactivation was then performed as described (Gerdes et al. 2013). Briefly, imaging was performed on a Cytell Slide Imaging System (GE Healthcare) at 20x magnification with exposure times determined for each antibody stain. Antibody reagents and staining sequence as described in Supplemental Table 1. Dye inactivation was accomplished with an alkaline peroxide solution, and background images were collected after each round of staining to ensure fluorophore inactivation.

### Image Processing

Following acquisition, images were processed as described (Gerdes et al. 2013; McKinley et al. 2017). Briefly, DAPI images for each round were registered to a common baseline, and autofluorescence in staining rounds was removed by subtracting the previous background image for each position. Images were then tiled for each TMA core.

### Cell Segmentation and Quantification Pipeline

All scripts to complete the segmentation pipeline are available at https://github.com/Coffey-Lab/CellSegmentation including a step-by-step overview of the process. Cell segmentation on each TMA core was conducted using a pipeline utilizing Matlab (R2018b) and Ilastik (1.3.2 or greater). Initialization of the Matlab wrapper function generated a number of container folders for each step in the segmentation results.

To facilitate pixel classification machine learning, tiff image stacks containing DAPI and autofluorescence removed images for all markers were generated for each image position after which the Matlab script was terminated. The tiff stacks were then manually annotated to generate epithelial/stroma, membrane/cytoplasm/nucleus, and stromal nuclei probability masks using a random forest pixel classification algorithm in Ilastik.

The Matlab script was re-initialized and binary masks for epithelium, nuclei, and membranes were generated. A watershed was used with nuclei as seed points to generate an initial segmentation, which was multiplied by the epithelial mask to only include epithelial cells. Subsequently, cells were resegmented using a novel algorithm if they contained greater than 10% internal membranes by area. A flowchart with pseudocode is shown in Figure S5. Briefly, line segments contained within an object are extended until they intersect with either the existing edge of the object or a separate extension within the object. The same method is also applied to cells expressing MUC2 in order to facilitate segmentation of goblet cells in intestinal tissues. Following re-segmentation, cells were assigned unique IDs, such that each contained within a cell is given the same pixel value.

Cell compartments were then defined within each cell. The nucleus was defined as the pixels contained within the cell object and the nuclear mask. Membranes were defined as the pixels contained within the cell object and within 5 pixels of the edge of the cell object that were not already defined as nuclear. The cytoplasm was defined as all other pixels in the cell object that were not already defined as nuclear or membranous. Each marker is then quantified over the entire cell and in each compartment by taking the median image intensity of the autofluorescence removed image. Additionally, the cell location (x- and y-coordinate of centroid) and areas of the entire cell object and subcompartmens were generated. These data were stored in CSV files.

For stromal cell segmentation, a watershed was used with stromal nuclei, defined from the machine learning pixel classification as seed points, was applied. This was then multiplied by a mask created from a 3 pixel dilation of the stromal nuclei mask to define the individual stromal cells. Marker quantification was then performed by calculating the median image intensity for each marker over the entire cell object. Cell location and area were also determined. These data were stored in CSV files.

### Cell Shape Analysis

A autoencoder neural network was then used to classify cell shapes from the segmented data. Similar methods have been developed to cluster non-biological data (Pimpaud 2019). For each imaging position, all segmented cells were binarized and re-sized to a 128 × 128 pixel matrix and aligned such that the major axes were all in the same orientation. The cell shape images were then saved as .mat files for each position. For autoencoder training, a random subset (20% default) of all cells were chosen and a neural network was run in Matlab using the Deep Learning Toolbox. Following training, all cells were encoded and the latent vectors for all cells was saved as a CSV file for later processing.

## Data Analysis

All analysis was performed in R. t-Stochastic Neighbor Embedding was conducted using the Rtsne package (Krijthe 2015) and all plots were generated using the ggplot2 package (Wickham 2016).

---

**Supplemental Figure 1.**
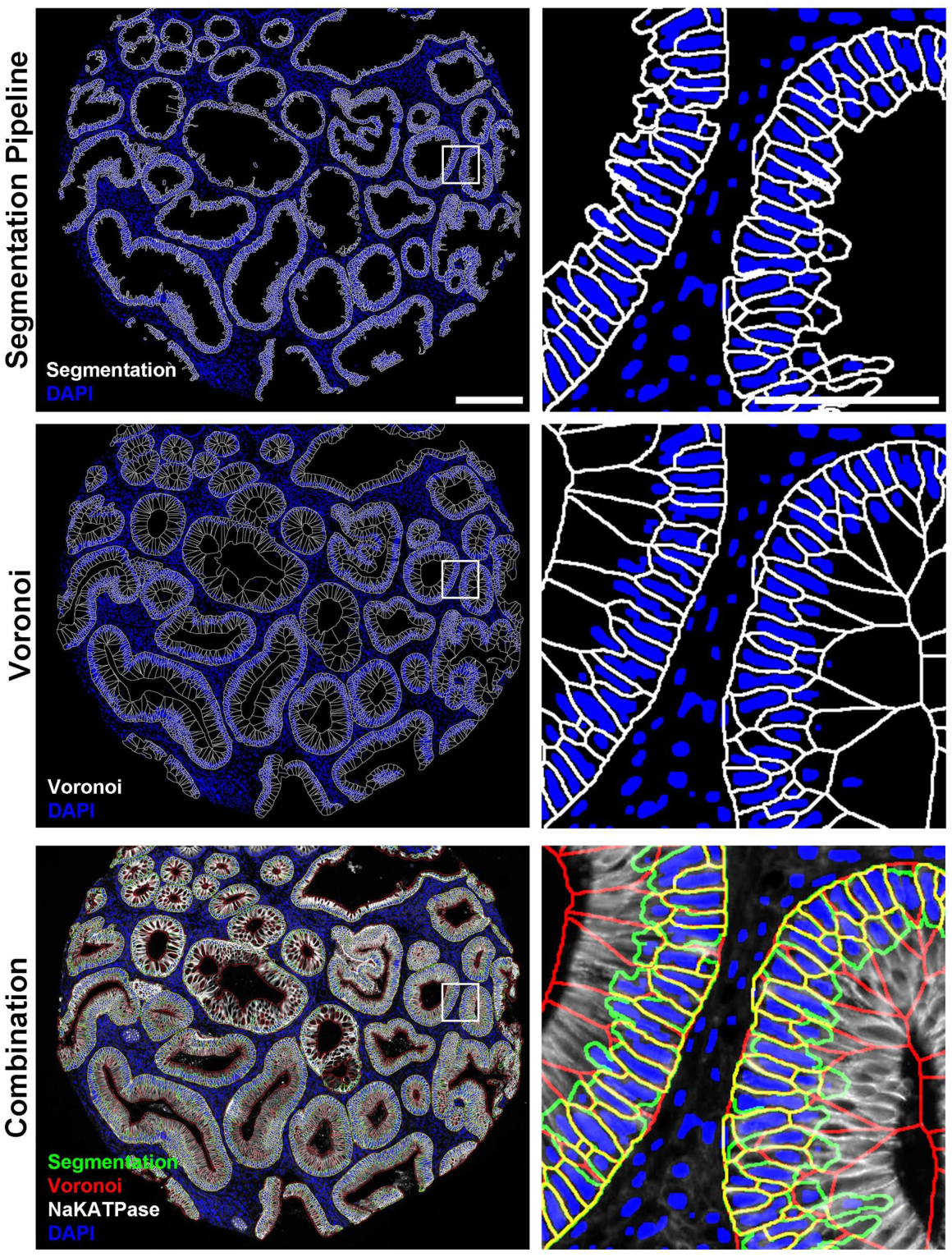
Comparison of segmentation pipeline and Voronoi. Segmentation results (white) with the nuclear mask are shown over an entire tissue microarray core (scale bar: 200 μm) and a zoomed region (scale bar: 50 μm). The same core is shown with voronoi segmentation using the nuclei as seed points. Overlaying both segmentation results demonstrates similar performance in regions of high nuclear density, but voronoi causes under-segmentation in regions where cell membranes exist, but where nuclei are not in the tissue section.

**Supplemental Figure 2.**
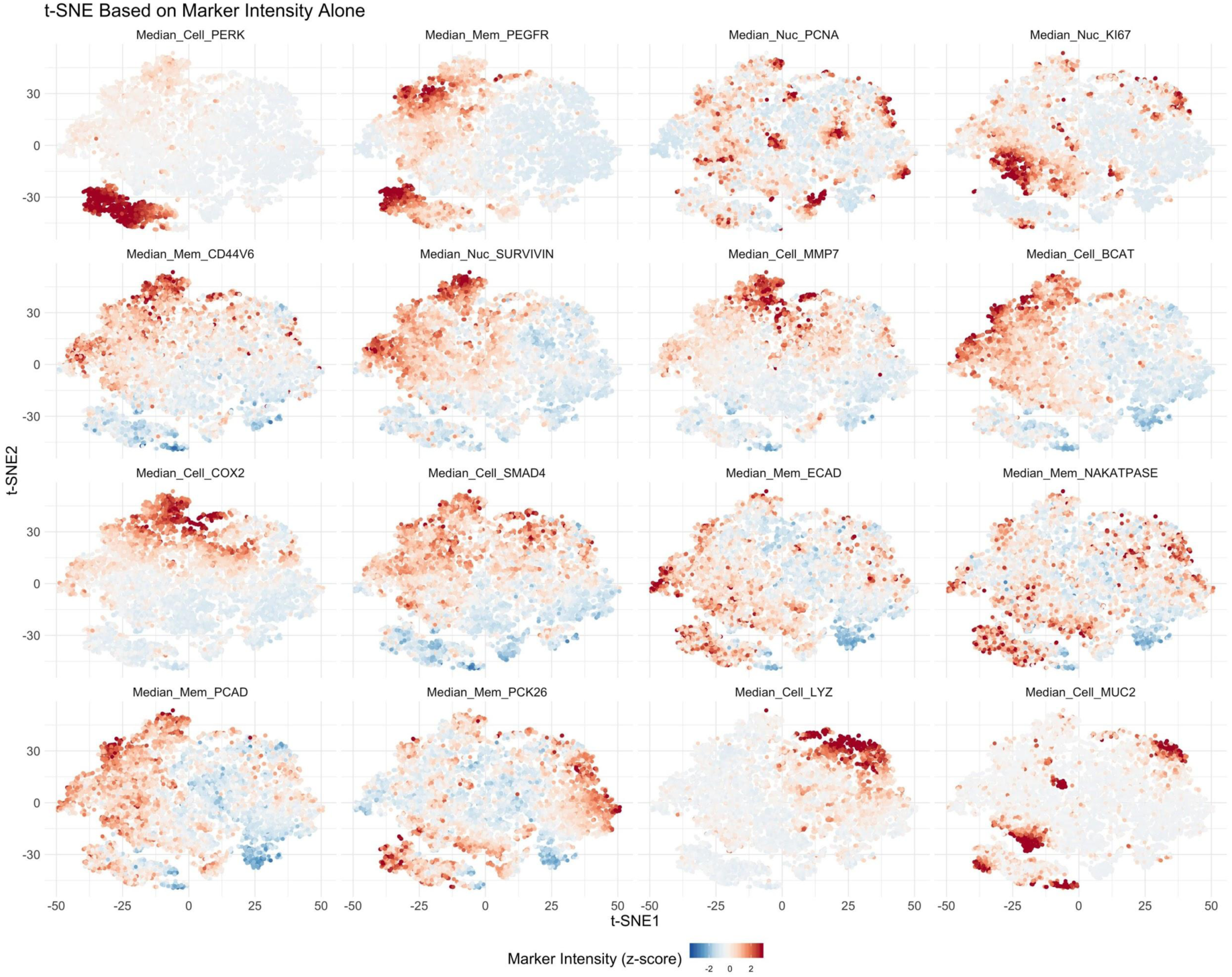
Marker intensity t-SNE. Plots are shown for each marker used for cell segmentation and t-SNE analysis using marker intensity only. A strong p-ERK-positive population can be identified in the lower left, while other populations are more diffuse.

**Supplemental Figure 3.**
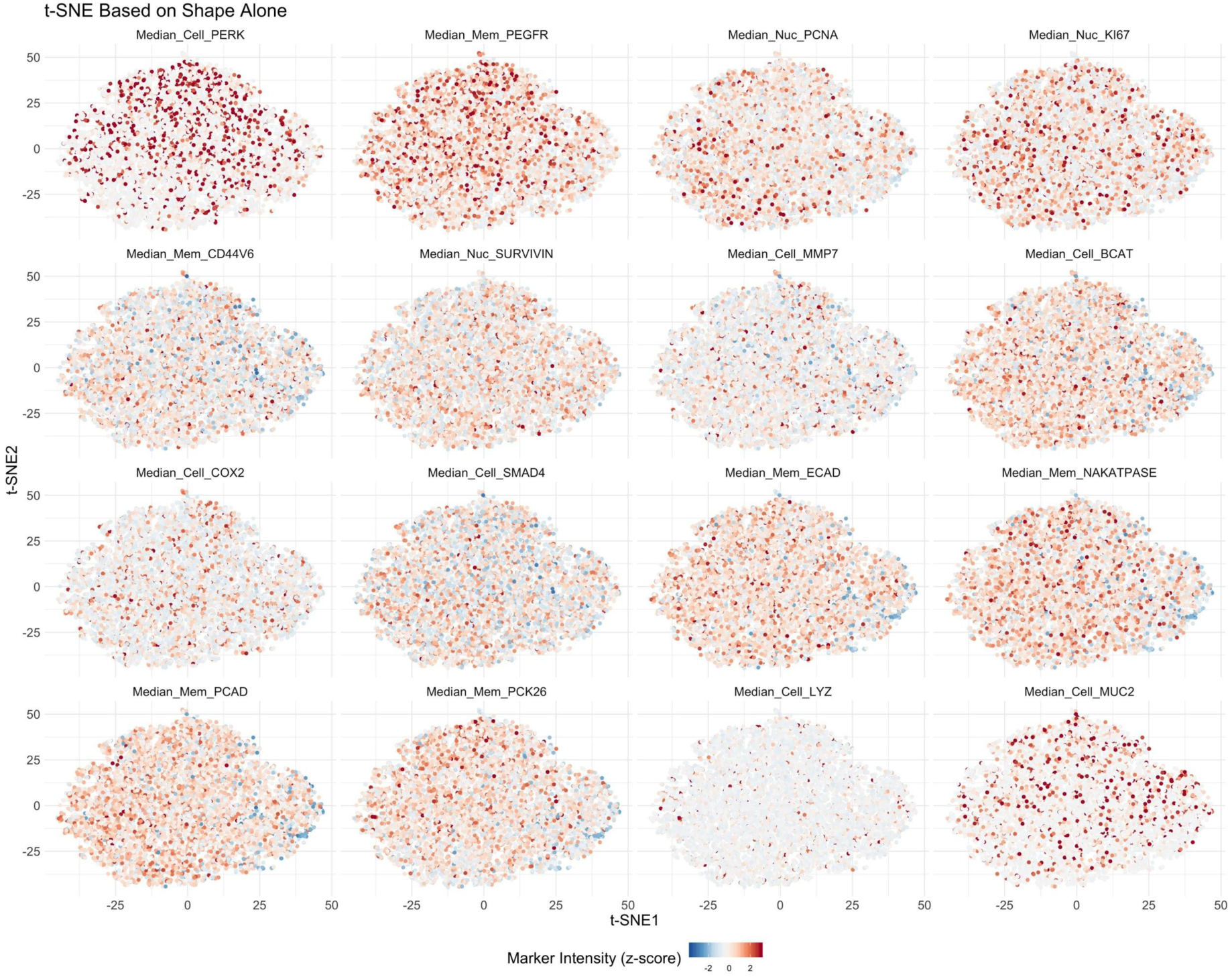
Shape t-SNE. Plots are shown for each marker used for cell segmentation and t-SNE analysis using only the cell shape latent vector as input. Unlike when staining intensity was used, no identifiable populations can be discerned with a seemingly random distribution for each marker demonstrating that shape is not reflecting marker expression.

**Supplemental Figure 4.**
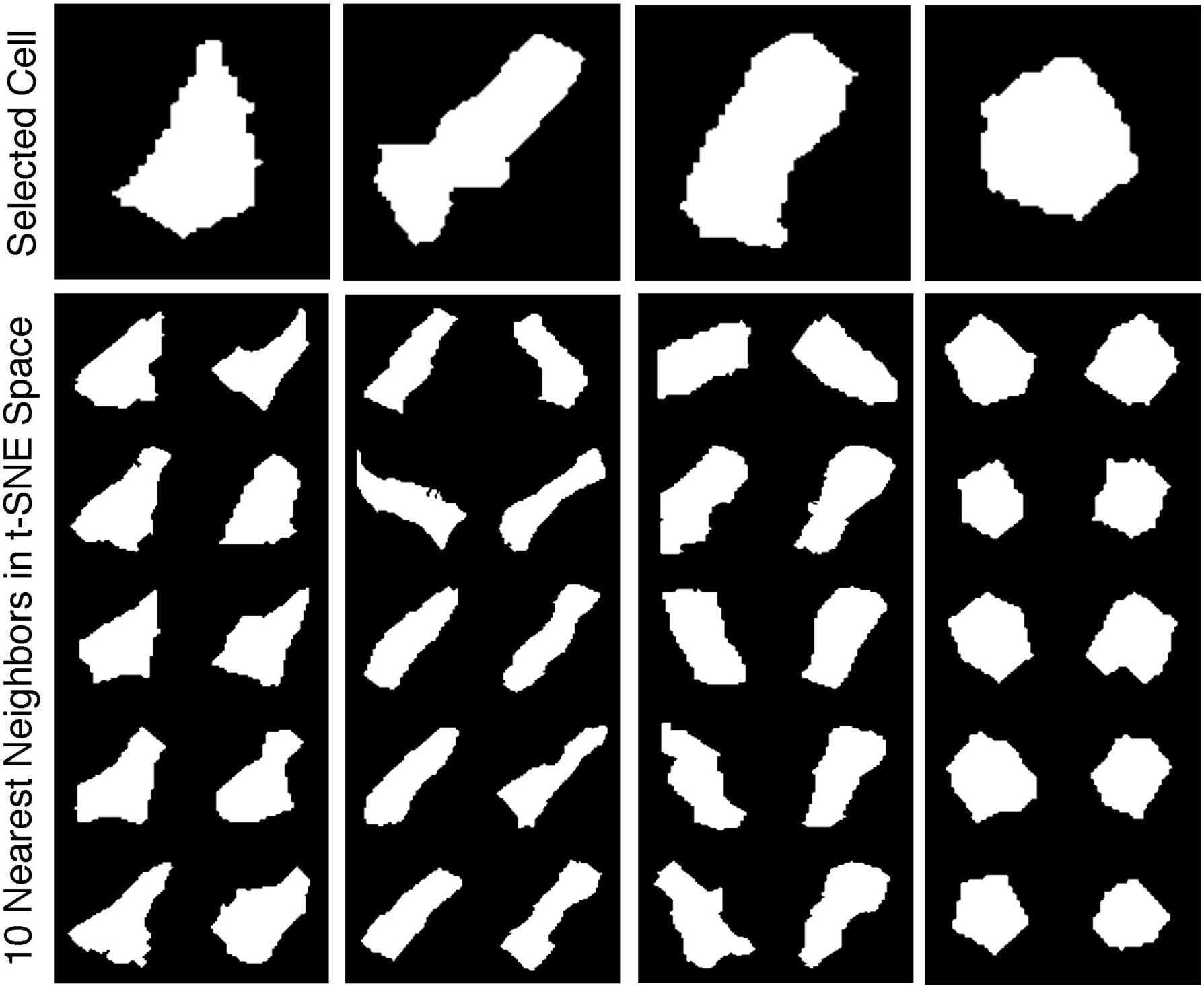
Cell shape similarity. Selected cells are compared to their 10 nearest t-SNE neighbors. The nearest neighbors are qualitatively similar in shape to each selected cell, demonstrating the ability of the autoencoder to characterize cell shapes.

**Supplemental Figure 5.**
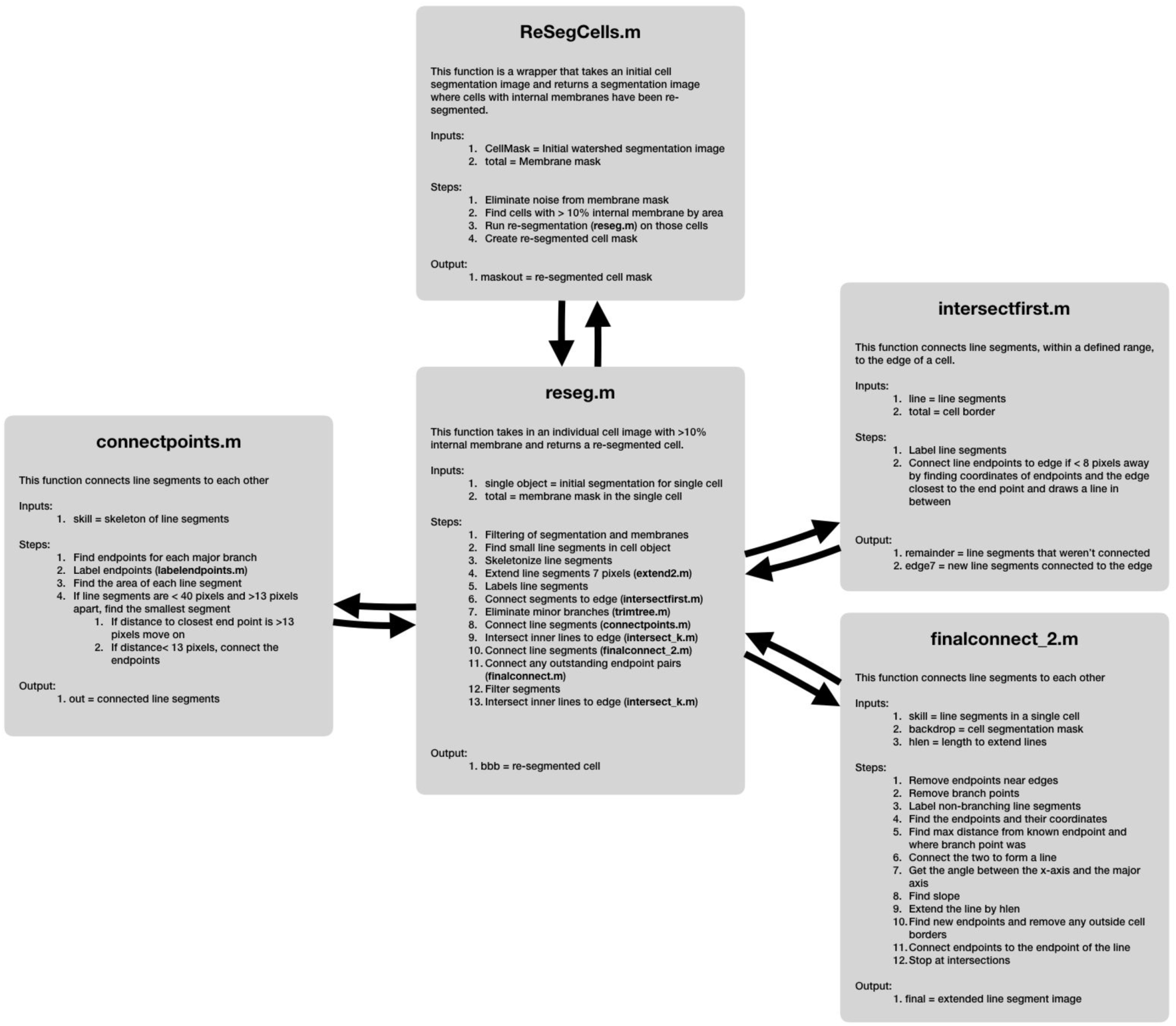
Cell re-segmentation algorithm. Major steps in the cell re-segmentation algorithm are shown with pseudocode.

## References

Al-Kofahi, Yousef, Alla Zaltsman, Robert Graves, Will Marshall, and Mirabela Rusu. 2018. “A Deep Learning-Based Algorithm for 2-D Cell Segmentation in Microscopy Images.” BMC Bioinformatics 19 (1): 365.

Angelo, Michael, Sean C. Bendall, Rachel Finck, Matthew B. Hale, Chuck Hitzman, Alexander D. Borowsky, Richard M. Levenson, et al. 2014. “Multiplexed Ion Beam Imaging of Human Breast Tumors.” Nature Medicine 20 (4): 436–42.

Baggett, Daniel, Masa-Aki Nakaya, Matthew McAuliffe, Terry P. Yamaguchi, and Stephen Lockett. 2005. “Whole Cell Segmentation in Solid Tissue Sections.” Cytometry. Part A: The Journal of the International Society for Analytical Cytology 67 (2): 137–43.

Bendall, Sean C., Erin F. Simonds, Peng Qiu, El-Ad D. Amir, Peter O. Krutzik, Rachel Finck, Robert V. Bruggner, et al. 2011. “Single-Cell Mass Cytometry of Differential Immune and Drug Responses across a Human Hematopoietic Continuum.” Science 332 (6030): 687–96.

Bradford, Jolene A., Gayle Buller, Michael Suter, Michael Ignatius, and Joseph M. Beechem. 2004. “Fluorescence-Intensity Multiplexing: Simultaneous Seven-Marker, Two-Color Immunophenotyping Using Flow Cytometry.” Cytometry. Part A: The Journal of the International Society for Analytical Cytology 61 (2): 142–52.

Gerdes, Michael J., Christopher J. Sevinsky, Anup Sood, Sudeshna Adak, Musodiq O. Bello, Alexander Bordwell, Ali Can, et al. 2013. “Highly Multiplexed Single-Cell Analysis of Formalin-Fixed, Paraffin-Embedded Cancer Tissue.” Proceedings of the National Academy of Sciences 110 (29): 11982–87.

Giesen, Charlotte, Hao A. O. Wang, Denis Schapiro, Nevena Zivanovic, Andrea Jacobs, Bodo Hattendorf, Peter J. Schüffler, et al. 2014. “Highly Multiplexed Imaging of Tumor Tissues with Subcellular Resolution by Mass Cytometry.” Nature Methods 11 (4): 417–22.

Goltsev, Yury, Nikolay Samusik, Julia Kennedy-Darling, Salil Bhate, Matthew Hale, Gustavo Vazquez, Sarah Black, and Garry P. Nolan. 2018. “Deep Profiling of Mouse Splenic Architecture with CODEX Multiplexed Imaging.” Cell 174 (4): 968–81.e15.

Herring, Charles A., Amrita Banerjee, Eliot T. McKinley, Alan J. Simmons, Jie Ping, Joseph T. Roland, Jeffrey L. Franklin, et al. 2018. “Unsupervised Trajectory Analysis of Single-Cell RNA-Seq and Imaging Data Reveals Alternative Tuft Cell Origins in the Gut.” Cell Systems 6 (1): 37–51.e9.

Krijthe, Jesse H. 2015. “Rtsne: T-Distributed Stochastic Neighbor Embedding Using Barnes-Hut Implementation.” R Package Version 0. 13, URL https://github.com/jkrijthe/Rtsne.

Lin, Jia-Ren, Benjamin Izar, Shu Wang, Clarence Yapp, Shaolin Mei, Parin M. Shah, Sandro Santagata, and Peter K. Sorger. 2018. “Highly Multiplexed Immunofluorescence Imaging of Human Tissues and Tumors Using T-CyCIF and Conventional Optical Microscopes.” eLife 7 (July). https://doi.org/10.7554/eLife.31657.

Maaten, Laurens van der, and Geoffrey Hinton. 2008. “Visualizing Data Using T-SNE.” Journal of Machine Learning Research: JMLR 9 (Nov): 2579–2605.

McKinley, Eliot T., Yunxia Sui, Yousef Al-Kofahi, Bryan A. Millis, Matthew J. Tyska, Joseph T. Roland, Alberto Santamaria-Pang, et al. 2017. “Optimized Multiplex Immunofluorescence Single-Cell Analysis Reveals Tuft Cell Heterogeneity.” JCI Insight 2 (11). https://doi.org/10.1172/jci.insight.93487.

Möller, Birgit, Yvonne Poeschl, Romina Plötner, and Katharina Bürstenbinder. 2017. “PaCeQuant: A Tool for High-Throughput Quantification of Pavement Cell Shape Characteristics.” Plant Physiology 175 (3): 998–1017.

Pimpaud, Benoit. 2019. “After Raw Stats: Exploring Possession Styles with Data Embeddings.” Medium. Towards Data Science. June 5, 2019. https://towardsdatascience.com/after-raw-stats-exploring-possession-styles-with-data-embeddings-d3ebef718abf.

Santamaria-Pang, A., J. Rittscher, M. Gerdes, and D. Padfield. 2015. “Cell Segmentation and Classification by Hierarchical Supervised Shape Ranking.” In 2015 IEEE 12th International Symposium on Biomedical Imaging (ISBI), 1296–99.

Schmitt, Oliver, and Maria Hasse. 2009. “Morphological Multiscale Decomposition of Connected Regions with Emphasis on Cell Clusters.” Computer Vision and Image Understanding: CVIU 113 (2): 188–201.

Schüffler, Peter J., Denis Schapiro, Charlotte Giesen, Hao A. O. Wang, Bernd Bodenmiller, and Joachim M. Buhmann. 2015. “Automatic Single Cell Segmentation on Highly Multiplexed Tissue Images.” Cytometry. Part A: The Journal of the International Society for Analytical Cytology 87 (10): 936–42.

Shrubsole, Martha J., Huiyun Wu, Reid M. Ness, Yu Shyr, Walter E. Smalley, and Wei Zheng. 2008. “Alcohol Drinking, Cigarette Smoking, and Risk of Colorectal Adenomatous and Hyperplastic Polyps.” American Journal of Epidemiology 167 (9): 1050–58.

Sommer, C., C. Straehle, U. Köthe, and F. A. Hamprecht. 2011. “Ilastik: Interactive Learning and Segmentation Toolkit.” In 2011 IEEE International Symposium on Biomedical Imaging: From Nano to Macro, 230–33.

Tang, Fuchou, Catalin Barbacioru, Yangzhou Wang, Ellen Nordman, Clarence Lee, Nanlan Xu, Xiaohui Wang, et al. 2009. “mRNA-Seq Whole-Transcriptome Analysis of a Single Cell.” Nature Methods 6 (5): 377–82.

Wickham, Hadley. 2016. ggplot2: Elegant Graphics for Data Analysis. Springer.

Yin, Z., R. Bise, M. Chen, and T. Kanade. 2010. “Cell Segmentation in Microscopy Imagery Using a Bag of Local Bayesian Classifiers.” In 2010 IEEE International Symposium on Biomedical Imaging: From Nano to Macro, 125–28.

Zimmer, Christophe, Elisabeth Labruyère, Vannary Meas-Yedid, Nancy Guillén, and Jean-Christophe Olivo-Marin. 2002. “Segmentation and Tracking of Migrating Cells in Videomicroscopy with Parametric Active Contours: A Tool for Cell-Based Drug Testing.” IEEE Transactions on Medical Imaging 21 (10): 1212–21.

